# Multi-centre laboratory study to determine discriminating concentrations for broflanilide and isocycloseram resistance monitoring in mosquitoes

**DOI:** 10.64898/2026.05.22.726487

**Authors:** Giorgio Praulins, Frank Mechan, Gemma Harvey, Basil D. Brooke, Vincent Corbel, Stéphane Duchon, Maria L. Kaiser, Sarah Moore, Ahmadi Bakari Mpelepele, Shüné V. Oliver, Himmat Singh, Jennifer Stevenson, Yvan G. Fotso Toguem, Vaishali Verma, Charles Wondji, Rosemary Susan Lees

**Author notes:** **Correspondence:** Rosemary Susan Lees.

## Abstract

In 2024–2025 a multi-centre study involving seven international laboratories was conducted with the support of the World Health Organization (WHO). The aim of the study was to establish and validate discriminating concentrations (DCs) in WHO bottle bioassays for monitoring susceptibility to broflanilide and isocycloseram in *Anopheles gambiae s*.*s*., *An. funestus, An. stephensi* and *Aedes aegypti*. The following values are recommended for adoption as DCs for broflanilide: 10 µg/bottle for *An. gambiae* and *Ae. aegypti*, 15 µg/bottle for *An. funestus*, and 25 µg/bottle for *An. stephensi*. The recommended DCs for isocycloseram are 15 µg/bottle for *Ae. aegypti*, 30 µg/bottle for *An. gambiae*, 50 µg/bottle for *An. stephensi*, and 60 µg/bottle for *An. funestu*s. Based on the experiences of conducting this study, which represents the application of a generic protocol for establishing discriminating concentrations produced by the WHO, technical recommendations are made on the generation and analysis of DC data for insecticides in future.

## 2. Introduction

The World Health Organization recommends that programmes routinely monitor target mosquito populations for susceptibility to insecticides to guide decisions about which insecticide-based vector control interventions to deploy (1). Field populations are regularly, typically annually, tested using bioassays measuring mortality on exposure to a known standard concentration of a given insecticide – known as the discriminating concentration (DC - sometimes referred to as the diagnostic concentration). Some insecticides are tested using treated filter papers in a WHO tube assay (2), and others in a treated Wheaton bottle via the WHO bottle bioassay (2). WHO has traditionally defined its discriminating concentrations (DCs) as twice the lowest concentration that gave systematically 100% mortality after 60 min exposure and a holding period of 24 hours using an insecticide susceptible strain. The DC is calculated as twice the LC_99_ value (concentration at which 99% of individuals are killed) as determined by log-probit statistical model against a susceptible strain (3). The concept of DCs has clear advantages in terms of the cost and efficiency of testing and has been widely adopted for the purposes of monitoring insecticide resistance in mosquitoes and other disease vectors.

In 2017-2021, WHO conducted a multi-centre study to establish and validate DCs of insecticides to enable the testing of resistance to insecticides in the main *Anopheles* and *Aedes* spp. mosquito vectors of human diseases (4). In 2022, the WHO published a methodology and a list of 12 new DCs to monitor insecticide resistance in *Culex quinquefasciatus* and *Cx. tarsalis* (5). Since these multi-centre studies, two new insecticides, both in class 30 of the Insecticide Resistance Action Committee (IRAC) classification have been included in products which have been listed as prequalified by WHO, broflanilide and isocycloseram (6). For the purpose of establishing and validating discriminating concentrations for these new insecticides, a multi-centre study was therefore conducted in 2024-5 to confirm and optimise where necessary the testing methods for performing dose response experiments with broflanilide and isocycloseram. DCs of each compound were established using the data analysis method developed by Mara Kont (7) and validated against well-characterised susceptible colonies of four mosquito species (*An. gambiae, An. funestus, An. stephensi*, and *Ae. aegypti*) with results generated from seven testing centres. Based on the experiences at the testing centres and with analysing the data in order to propose DCs, the WHO’s generic protocol for establishing DCs was further validated, and recommendations on best practice were formulated.

## 3 Materials and Methods

### 3.1 Participating institutions

Seven internationally recognised laboratories with high entomological capacity participated in the study, which was coordinated by the Liverpool School of Tropical Medicine (LSTM) in collaboration with WHO. These were: LSTM, Institut de Recherche pour le Développement (IRD), London School of Hygiene & Tropical Medicine (LSHTM), Centre for Research in Infectious Diseases (CRID), ICMR–National Institute of Malaria Research (NIMR), National Institute for Communicable Diseases (NICD), and Ifakara Health Institute (IHI). All participating institutions maintained well-characterised susceptible mosquito colonies with adequate facilities and technical capacity to conduct laboratory testing. These institutions had previously participated in WHO multi-centre trials or demonstrated strong capacity for insecticide testing.

### 3.2 Study design

The study followed a step-by-step approach (Figure 1) previously used (4,5,8) and now formalised by WHO as a generic protocol for studies to determine discriminating concentrations (3) with the aim of establishing broflanilide and isocycloseram DCs for inclusion in the WHO Manual for monitoring insecticide resistance in mosquito vectors and selecting appropriate interventions (1).

**Figure 1.**
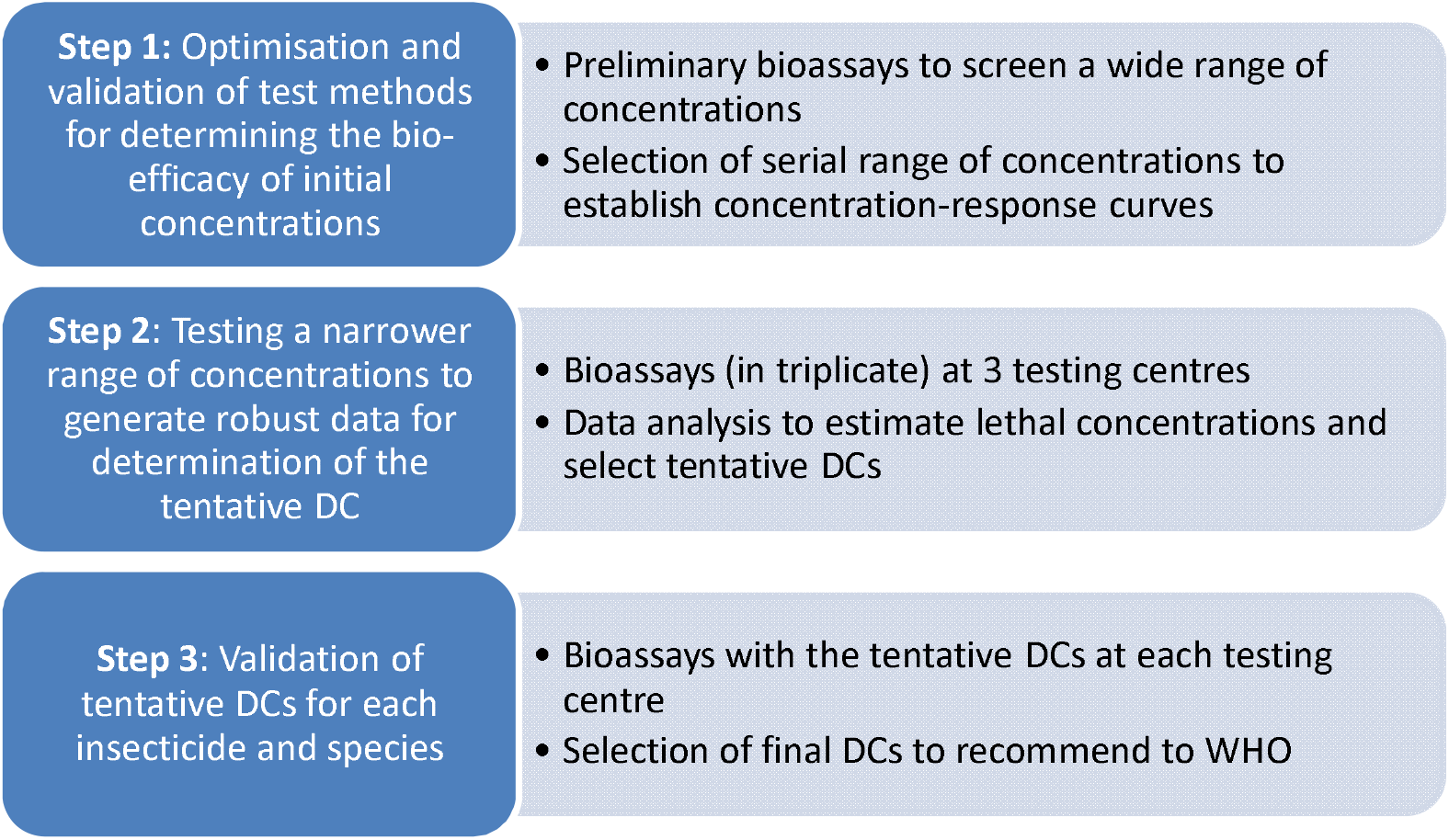
Schematic representation of the study design to determine discriminating concentrations of insecticides for monitoring resistance in mosquitoes. Adapted from *(3)*.

#### Step 1: Preliminary bottle bioassays for screening a range of serial concentrations of insecticides to obtain 0–100% mortality in mosquitoes

About 10– 12 serial concentrations were required to provide a range of mortalities (from 0 to 100%) for each mosquito species and compound. Susceptible mosquitoes were exposed to serial concentrations of each insecticide for 1h, at one replicate each of 50 mosquitoes per concentration and one replicate of 50 mosquitoes as an acetone-only control.

#### Step 2. Bottle bioassays to establish LC_99_ and LC_100_ and propose tentative discriminating concentrations

For Step 2 testing, each centre was expected to complete three validated replicates for each concentration of each compound against each of their representative species (Table 1). The overall aim was to achieve a complete dataset of three replicates from each of three centres for every compound–species pair. Testing centres tested a range of serial concentrations pre-selected by LSTM based on the results of Step 1 testing. At least six test concentrations and a control were tested to generate “concentration-response” curves. For a Step 2 test to be considered valid, it was necessary to select at least two concentrations killing <50% mosquitoes, one concentration killing around 50%, two concentrations killing >50% mosquitoes and one concentration killing »100% of mosquitoes. Each bioassay was performed in triplicate for a given species and mosquito strain (with at least 100 mosquitoes per concentration) using different batches of mosquitoes on different days. The impregnated bottles were not used more than once.

**Table 1.**
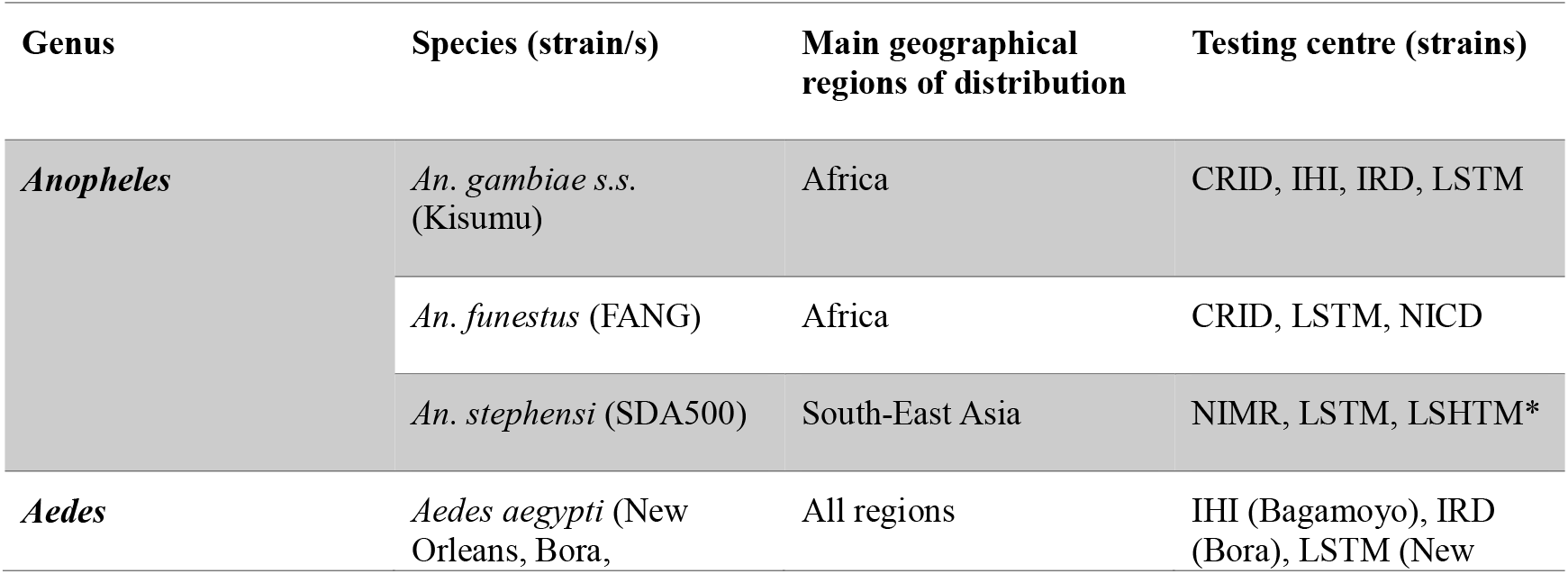
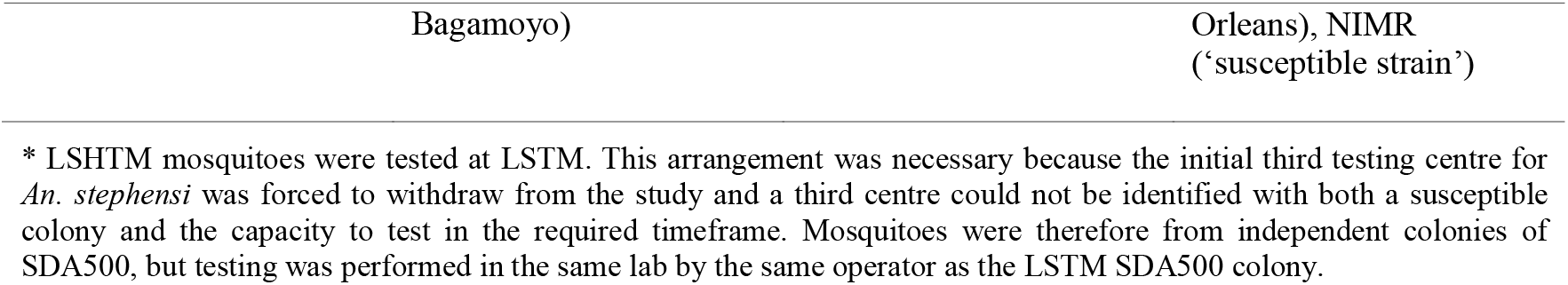
Susceptible mosquito species and strains used for the study, and the testing centres that tested each.

#### Step 3. Multi-centre validation of tentative discriminating concentrations against all mosquito species

Tentative DCs (TDCs) were calculated based on the data from Step 1, in discussion with testing centres and WHO representatives. The TDC of a given insecticide was tested by three different laboratories against the same well-characterized susceptible strains in WHO bottle bioassays, recording 24-hour mortality. Bioassays included one replicate for each compound and mosquito species, with at least 100 mosquitoes tested per tentative DC to validate whether 100% of susceptible strains tested were killed.

### 3.3 Mosquitoes

Four mosquito vector species (*An. gambiae, An. funestus, An. stephensi*, and *Ae. aegypti*) were included in the study based on three selection criteria: i. species of major epidemiological importance in malaria or dengue transmission, ii. representation of different geographical regions (Africa, Asia, and other relevant endemic areas), and iii. availability of well-characterized, fully susceptible colonies maintained in at least three participating laboratories, ensuring cross-validation of results across multiple sites (see Table 1). The mosquito species and susceptible strains tested at each centre were selected according to local colony availability and laboratory capacity.

### 3.4 WHO bottle bioassays

Given the chemical properties and modes of action of broflanilide and isocycloseram, and the experience of their manufacturers who were consulted ahead of the study, the WHO bottle bioassay (2,7), with the addition of a surfactant (rapeseed oil methyl ester, MERO) was selected as the most appropriate testing methodology. Broflanilide was provided to LSTM for the study by Mitsui Chemicals Crop & Life Solutions, Inc., and isocycloseram by Syngenta Crop Protection AG, and aliquots of each insecticide and of MERO was sent to each testing centre.

Bottles were coated with the required doses of insecticide and MERO surfactant at 1,500 ppm for *Aedes aegypti* and 800 ppm for *Anopheles* spp. according to the WHO bottle bioassay SOP (2). The bottle drying time (i.e. interval of time between coating bottles and testing) was fixed at 24h, and bottles were dried horizontally with their lids remaining open until the test started.

3-5 day old non-blood fed female mosquitoes were exposed in the treated bottles for 1 hour according to the WHO bottle bioassay SOP (2). Testing was performed and mosquitoes held until the 24h mortality scoring at 27° ± 2°C and 75 ± 10% relative humidity; all testing centres reported testing conditions within this range.

### 3.5 Data analysis

To ensure consistency and robustness in defining the DCs before progressing to the next step of validation, bioassay data for Step 2 were included for analysis only if the control mortality was below 20%, a minimum of six concentrations were tested for generation of concentration-response curves and to estimate LC values, the goodness of fit had to meet the threshold (P-value > 0.05), and the dataset had to include a concentration that killed 100% of mosquitoes (LC_100_), where available.

For each testing centre, data from three replicates were analysed separately to assess intra-laboratory variation. If no major differences were observed between replicates, the data were pooled to estimate LC values for that centre. Datasets from three independent laboratories were used to cross-validate results and determine a tentative DC. Concentration response curves were plotted for visualisation of the data. Concentration response data for each insecticide–species combination were analysed using the Bayesian binomial modelling framework developed for the WHO bottle bioassay multi-centre study. For each bioassay, the number of mosquitoes dead out of those tested was modelled using a five-parameter logistic regression on square-root transformed concentration scales. Models were run on four Markov chains for up to 10,000 iterations, and convergence was assessed using the Gelman– Rubin statistic (R□ < 1.01) with zero divergent iterations required. All models presented in Table 3 successfully met these convergence criteria.

The fitted curves and confidence intervals were used to assess consistency across test centres and species and to derive LC and tentative discriminating concentration estimates (7). The DC for each insecticide and each species was defined as twice the estimated LC_99_. If differences in LC_99_ were observed between centres, the highest LC_99_ value was selected. In cases where an LC_99_ could not be accurately estimated (e.g., due to insufficient data points or replicates), an observed LC_100_ was instead used to establish the DC.

For Step 3 the concentration-response bioassay data were analysed, and the concentration-response parameters were used to estimate the goodness of fit (with the P-value) and lethal concentrations (i.e., LC_100_ and LC_99_ with 95% confidence intervals) for each insecticide and mosquito species.

## 4 Results

### 4.1 Step 1. Preliminary bottle bioassays for screening a range of serial concentrations of insecticides to obtain 0–100% mortality in mosquitoes

Preliminary data generated at LSTM and IRD confirmed the suitability of a 24h drying time for bottles before testing, a one-hour exposure period, and mortality 24h post-exposure to be a suitable endpoint. The initial concentrations used by each centre for testing in Step 2 were established based on preliminary testing at LSTM.

### 4.2 Step 2. Bottle bioassays to establish LC_99_ and LC_100_ and propose tentative discriminating concentrations

In total, 62,673 *Aedes* and *Anopheles* mosquitoes were tested in WHO bottle assays treated with broflanilide and isocycloseram (Table 2).

**Table 2.** Total number of mosquitoes tested in WHO bottle assays against broflanilide and isocycloseram by species.

| Species | Broflanilide | Isocycloseram | Total |
| --- | --- | --- | --- |
| *Ae. aegypti* | 10,285 | 8,673 | 18,958 |
| *An. gambiae* | 10,139 | 9,249 | 19,388 |
| *An. funestus* | 6,948 | 5,126 | 12,074 |
| *An. stephensi* | 3,949 | 4,552 | 8,501 |
| *An. coluzzii* | 1,856 | 1,896 | 3,752 |
| **Total** | 33,177 | 29,496 | 62,673 |

#### 4.2.1 Broflanilide

Where data sets were suitable, concentration response curves were plotted for each insecticide-species combination by testing centre; where data could not be fitted by the model, no curve was generated. The fitted curves and confidence intervals were used to assess consistency across test centres and species and to derive LC and tentative discriminating concentration estimates.

LSTM carried out a total of two replicates with An. gambiae, tested across six concentrations ranging from 0.05 to 5 µg/bottle, IRD carried out a total of three replicates tested across eight concentrations ranging from 0.08 to 1 µg/bottle, and IHI carried out a total of eight replicates tested across nine concentrations ranging from 0.25 to 25 µg/bottle (Figure 2). CRID tested three replicates across six concentrations however data could not be fitted by the model, so no curve was generated for that site.

**Figure 2.**
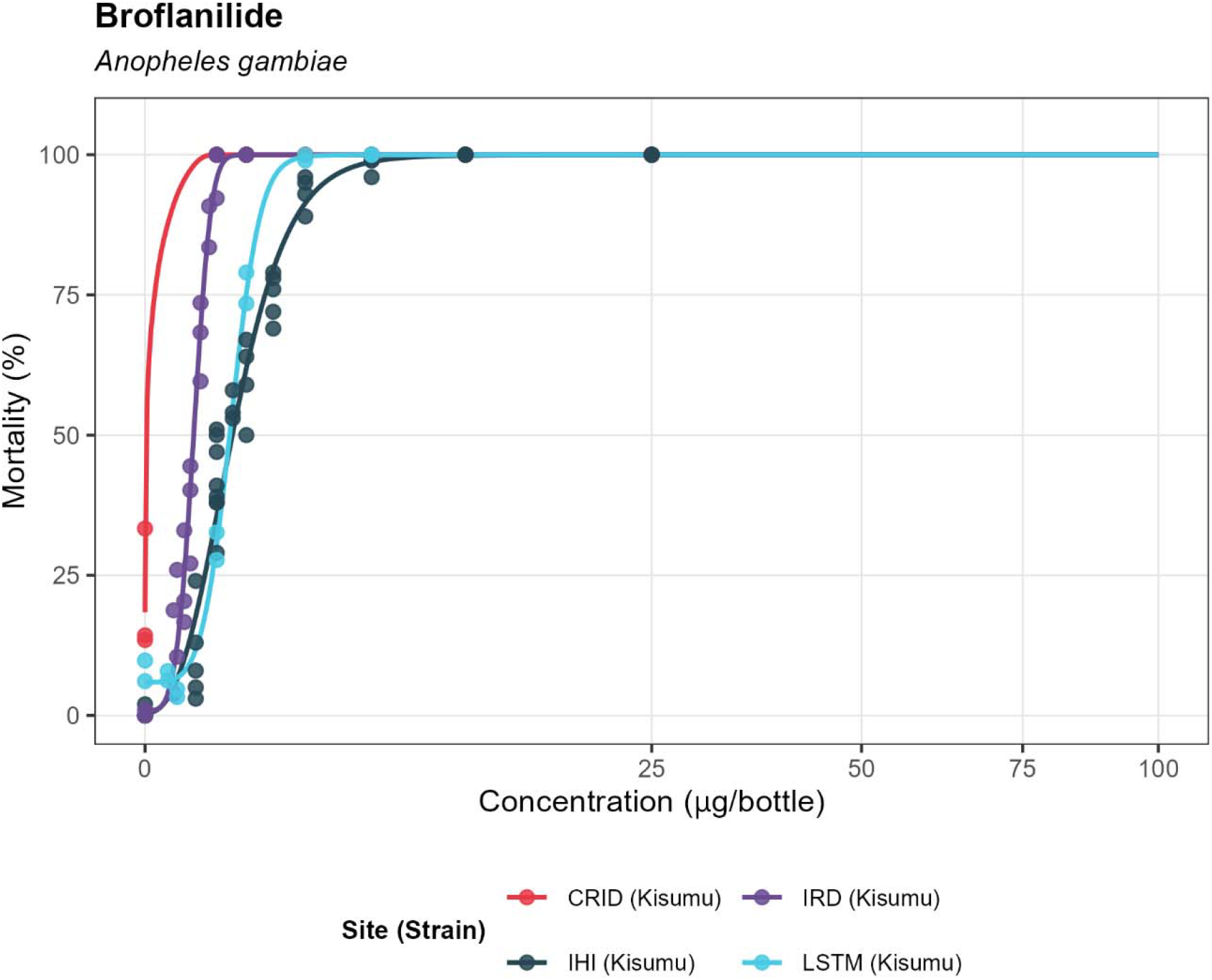
*Anopheles gambiae* mosquito mortality in bottle bioassays with broflanilide-coated bottles after a 24hr holding period at different test centres (test conditions: 1h exposure, 24hr bottle drying time).

Two centres generated data for *An. funestus* that could be fitted to the model; LSTM carried out six replicates tested across eight concentrations ranging from 0.1 to 25 µg/bottle, NICD carried out three replicates tested across eight concentrations ranging from 0.05 to 25 µg/bottle (Figure 3). CRID carried out a total of five replicates tested across eight concentrations ranging from 0.005 to 25 µg/bottle, but the data could not be fitted to the model.

**Figure 3.**
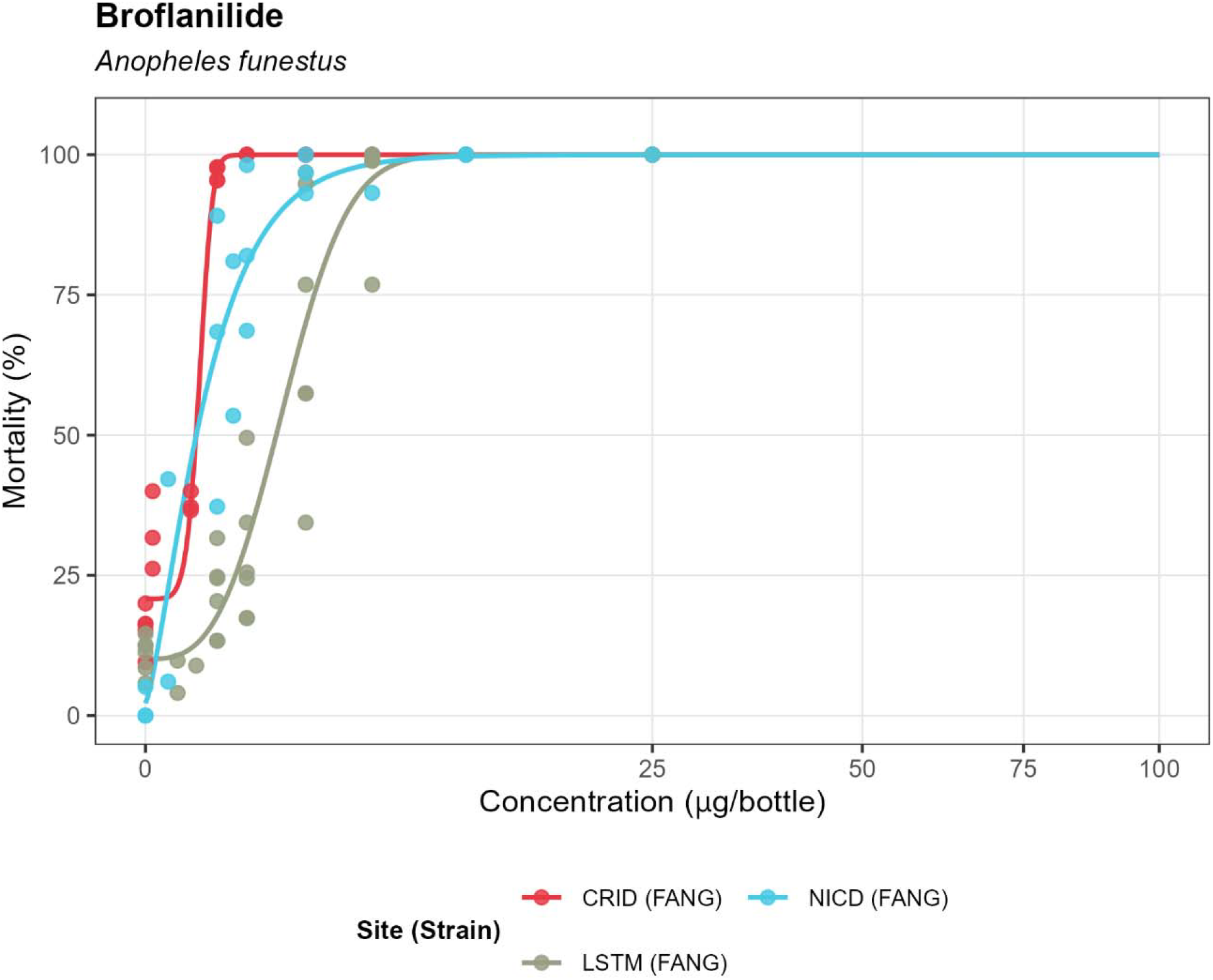
*Anopheles funestus* mosquito mortality in bottle bioassays with broflanilide-coated bottles after a 24hr holding period at different test centres (test conditions: 1h exposure, 24hr bottle drying time).

With *An. stephensi*, LSTM carried out two replicates tested across six concentrations ranging from 0.1 to 10 µg/bottle for the LSTM colony of SDA500, as well as two replicates tested across six concentrations ranging from 0.0005 to 5 µg/bottle for the LSHTM colony. NIMR carried out three replicates tested across six concentrations ranging from 0.5 to 25 µg/bottle with their susceptible strain (Figure 4).

**Figure 4.**
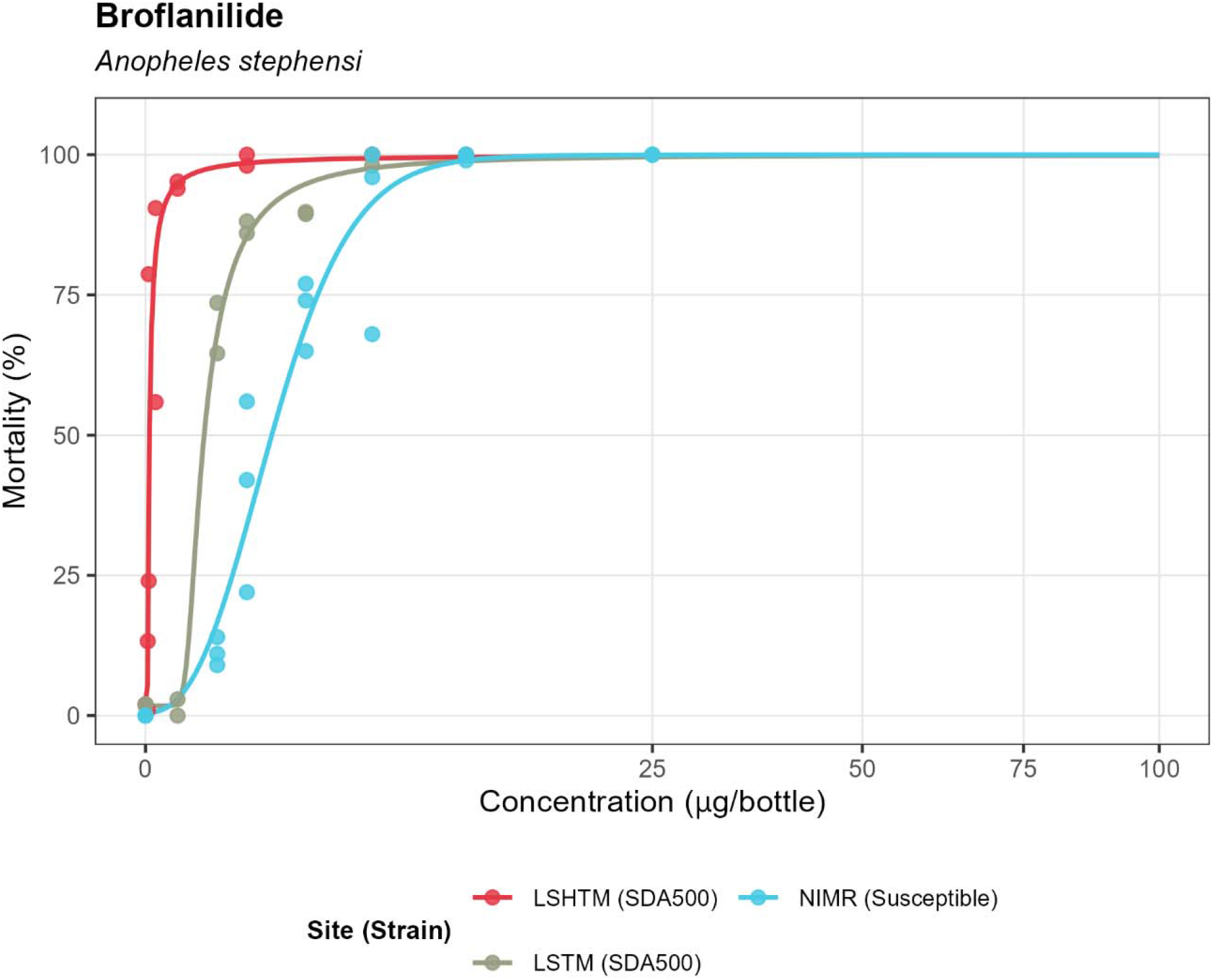
*Anopheles stephensi* mosquito mortality in bottle bioassays with broflanilide-coated bottles after a 24hr holding period at different test centres (test conditions: 1h exposure, 24hr bottle drying time).

Four centres generated data for *Aedes aegypti*. IHI carried out seven replicates tested across nine concentrations ranging from 0.25 to 25 µg/bottle, NIMR carried out three replicates tested across six concentrations ranging from 0.5 to 25 µg/bottle, LSTM carried out two replicates tested across six concentrations ranging from 0.05 to 5 µg/bottle and IRD carried out three replicates tested across eight concentrations ranging from 0.1 to 1.5 µg/bottle (Figure 5).

**Figure 5.**
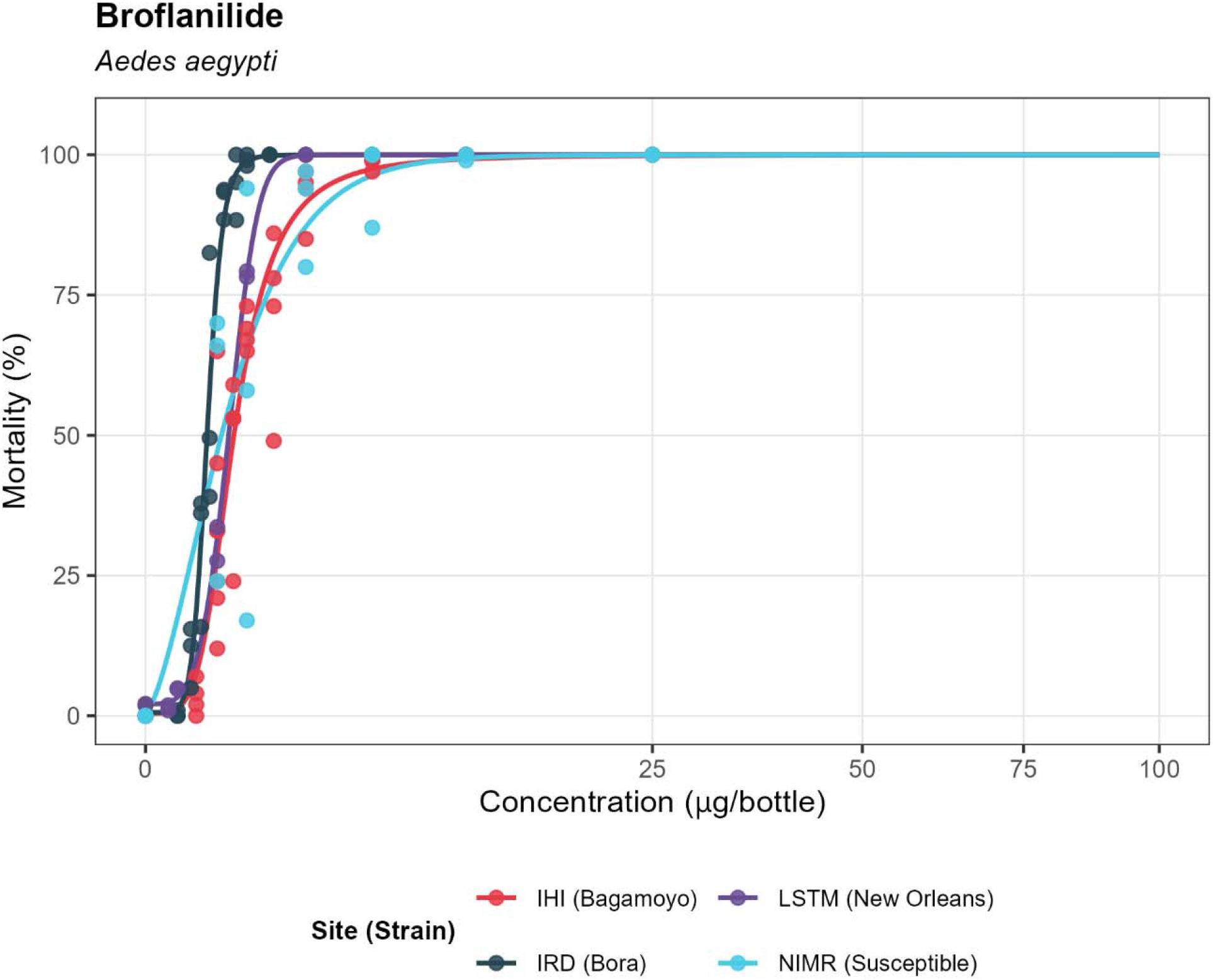
*Aedes aegypti* mosquito mortality in bottle bioassays with broflanilide-coated bottles after a 24hr holding period at different test centres (test conditions: 1h exposure, 24hr bottle drying time).

#### 4.2.2 Isocycloseram

With An. gambiae, IHI carried out six replicates with isocycloseram tested across nine concentrations ranging from 0.25 to 25 µg/bottle, IRD carried out three replicates tested across eight concentrations ranging from 0.05 to 1 µg/bottle and LSTM carried out two replicates tested across six concentrations ranging from 0.5 to 10 µg/bottle (Figure 6). CRID carried out three replicates tested across six concentrations ranging from 1 to 25 µg/bottle, but data could not be fitted by the model.

**Figure 6.**
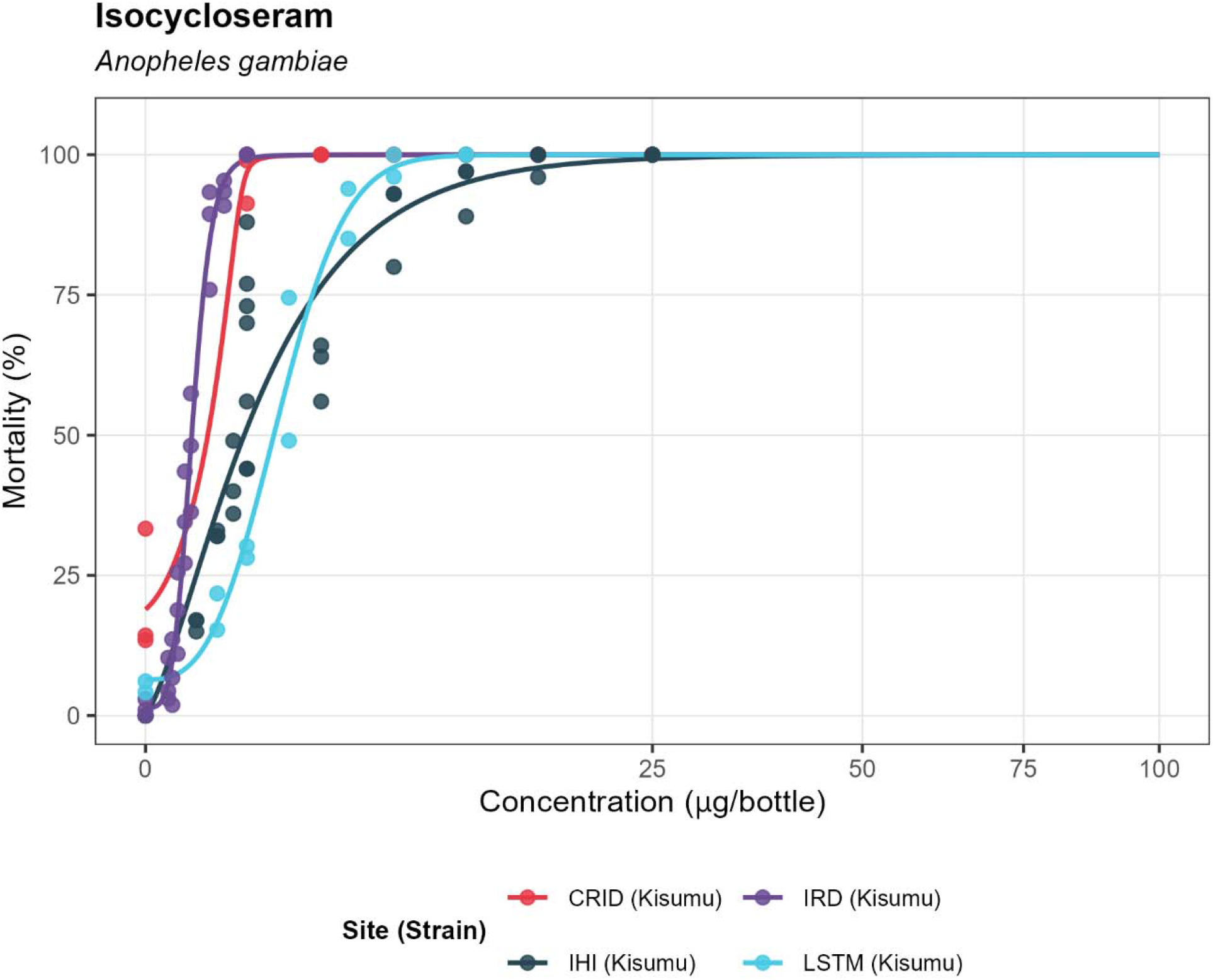
*Anopheles gambiae* mosquito mortality in bottle bioassays with isocycloseram-coated bottles after a 24hr holding period at different test centres (test conditions: 1h exposure, 24hr bottle drying time).

For *An. funestus*, NICD carried out two replicates tested across six concentrations ranging from 1 to 25 µg/bottle, CRID carried out four replicates tested across eight concentrations ranging from 0.25 to 25 µg/bottle and LSTM carried out five replicates tested across nine concentrations ranging from 0.35 to 100 µg/bottle (Figure 7).

**Figure 7.**
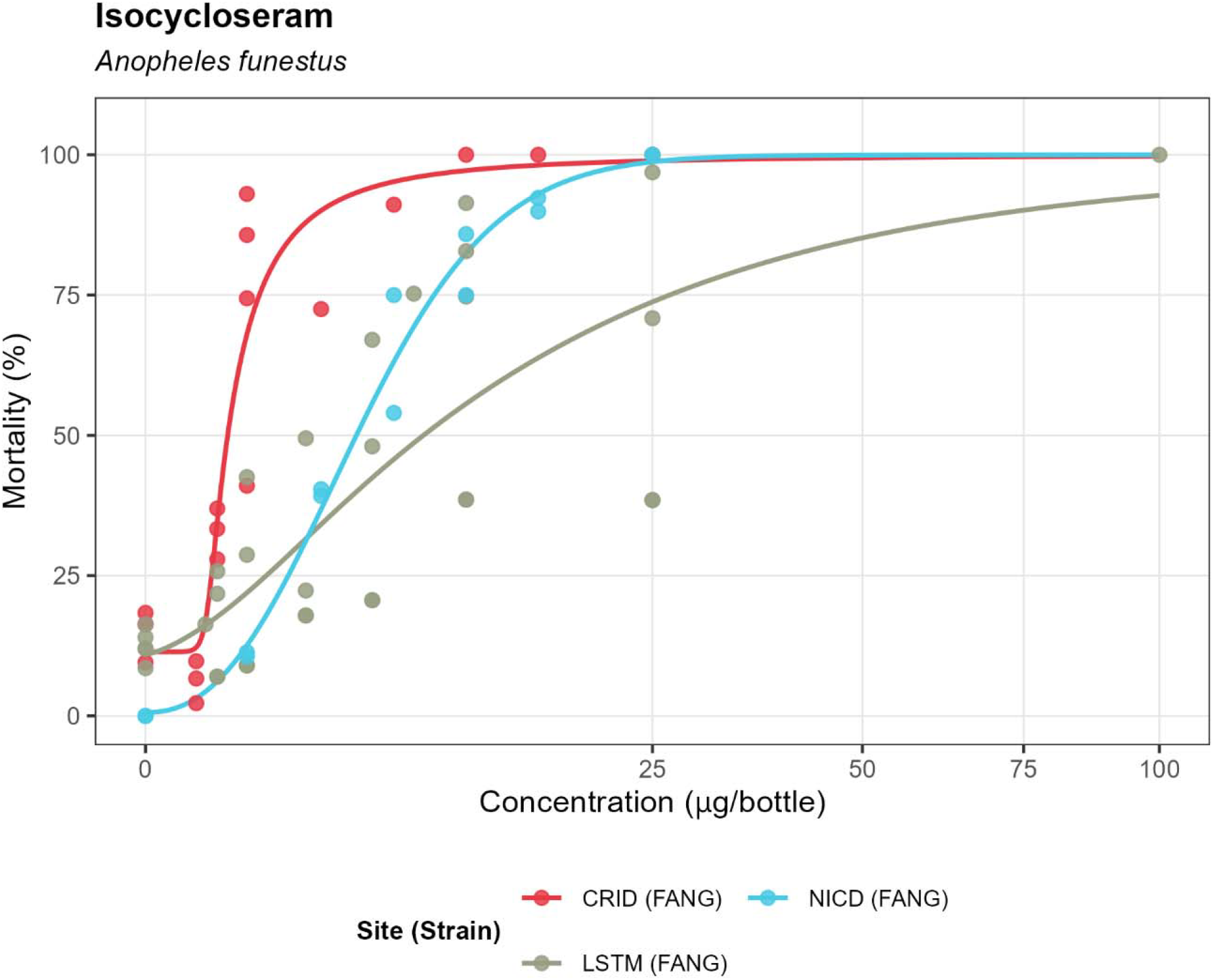
*Anopheles funestus* mosquito mortality in bottle bioassays with isocycloseram-coated bottles after a 24hr holding period at different test centres (test conditions: 1h exposure, 24hr bottle drying time).

NIMR carried out three replicates with *An. stephensi* tested across six concentrations ranging from 0.5 to 25 µg/bottle, LSTM carried out three replicates tested across six concentrations ranging from 0.5 to 25µg/bottle for the LSTM strain as well as two replicates tested across six concentrations ranging from 0.5 to 25 µg/bottle for the LSHTM strain (Figure 8). Data generated by LSTM with the LSHTM colony could not be fitted to the model.

**Figure 8.**
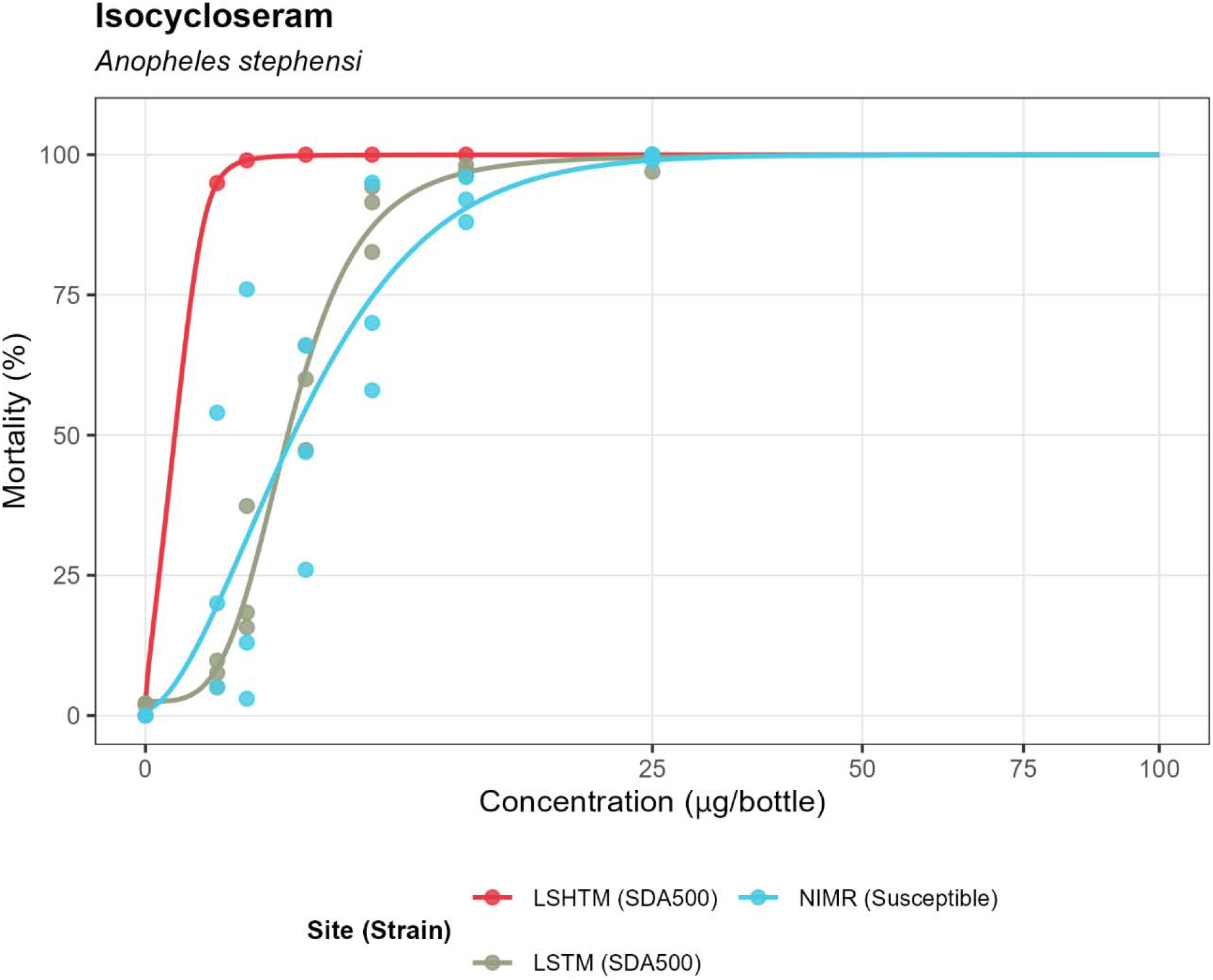
*Anopheles stephensi* mosquito mortality in bottle bioassays with isocycloseram-coated bottles after a 24hr holding period at different test centres (test conditions: 1h exposure, 24hr bottle drying time).

IHI carried out six replicates with *Ae. aegypti* tested across nine concentrations ranging from 0.25 to 25 µg/bottle, IRD carried out three replicates tested across six concentrations ranging from 0.3 to 2 µg/bottle, LSTM carried out two replicates tested across eight concentrations ranging from 0.05 to 10 µg/bottle and NIMR carried out three replicates tested across six concentrations ranging from 0.5 to 25 µg/bottle (Figure 9).

**Figure 9.**
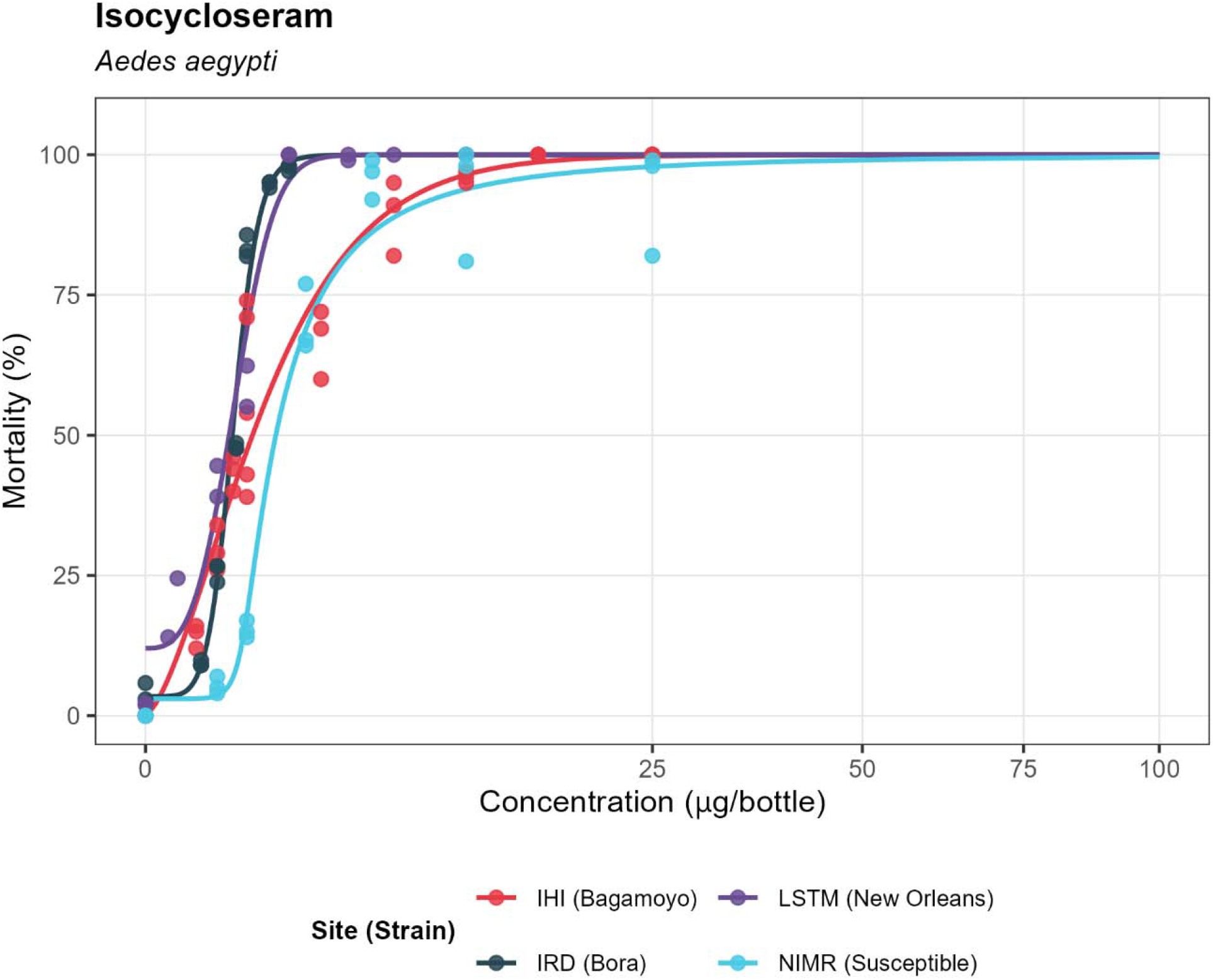
*Aedes aegypti* mosquito mortality in bottle bioassays with isocycloseram-coated bottles after a 24hr holding period at different test centres (test conditions: 1h exposure, 24hr bottle drying time).

For both broflanilide and isocycloseram, the fitted models produced steep slopes with narrow confidence intervals at higher mortality levels, indicating consistent concentration–response relationships. LC values were generally lower for *An. gambiae* and *Ae. aegypti* than *An. funestus* and *An. stephensi*, reflecting differences in baseline sensitivity. While some inter-laboratory variation was observed, particularly in LC□ □ estimates, the overall trends were coherent and suitable for establishing tentative discriminating concentrations based on pooled or representative site data according to the study protocol (Table 3).

**Table 3.** Summary of LC_99_ values, discriminating concentration (DC = 2 × LC_99_), and observed LC_100_ for broflanilide and isocycloseram across all participating centres, mosquito species, and strains in WHO bottle bioassays. Values are presented to one decimal place as median LC_99_ with lower and upper 95% confidence intervals, the corresponding DC, and the lowest tested concentration consistently giving 100% mortality (Observed LC_100_). Model fit was assessed following Kont et al. (2023) using the coefficient of determination (R^2^) and root mean square error (RMSE), calculated from a linear regression of observed against posterior median predicted mortality at each tested concentration; R^2^ values approaching 1.0 and RMSE values approaching 0% indicate good model fit. N/A indicates data not available or not determined. Results that did not meet convergence criteria (Gelman–Rubin statistic R□ < 1.01 with zero divergent iterations) are flagged; see footnote† for details.

| Insecticide | Site | Species | Strain | LC99<br>median | LC99<br>lower | LC99<br>upper | DC 2x<br>LC99 | Observed<br>LC100 | R<sup>2</sup> | RMSE | Converged |
| --- | --- | --- | --- | --- | --- | --- | --- | --- | --- | --- | --- |
| Broflanilide | IHI | Aedes<br>aegypti | Bagamoyo | 8.5 | 5.9 | 13.0 | 17.0 | 10.0 | 0.9 | 9.7 | Yes |
|  | IRD |  | Bora | 1.0 | 0.9 | 1.2 | 2.1 | 0.8 | 1.0 | 7.6 | Yes |
|  | LSTM |  | New Orleans | 1.8 | 1.5 | 2.3 | 3.6 | 2.5 | 1.0 | 1.5 | Yes |
|  | NIMR |  | Susceptible | 8.4 | 6.8 | 10.8 | 16.7 | 5.0 | 0.8 | 15.3 | Yes |
|  | CRID | Anopheles<br>coluzzii | N'gousso | 0.0 | 0.0 | 0.3 | 0.0 | 0.5 | 1.0 | 1.3 | No† |
|  | CRID | Anopheles<br>funestus | FANG | 0.6 | 0.5 | 0.7 | 1.2 | 1.0 | 1.0 | 5.3 | No† |
|  | LSTM |  | FANG | 6.8 | 6.1 | 7.6 | 13.5 | 5.0 | 0.9 | 10.4 | Yes |
|  | NICD |  | FANG | 6.2 | 4.8 | 8.6 | 12.3 | 2.5 | 0.9 | 12.1 | Yes |
|  | CRID | Anopheles<br>gambiae | Kisumu | 0.0 | 0.0 | 0.4 | 0.0 | 0.5 | 1.0 | 3.5 | No† |
|  | IHI |  | Kisumu | 5.3 | 4.4 | 7.3 | 10.6 | 5.0 | 1.0 | 5.9 | Yes |
|  | IRD |  | Kisumu | 0.6 | 0.6 | 0.7 | 1.2 | 0.5 | 1.0 | 5.9 | Yes |
|  | LSTM |  | Kisumu | 2.1 | 1.7 | 2.8 | 4.1 | 2.5 | 1.0 | 2.2 | Yes |
|  | LSHTM | Anopheles<br>stephensi | SDA500 | 2.1 | 0.9 | 5.8 | 4.2 | 1.0 | 0.9 | 13.9 | Yes |
|  | LSTM |  | SDA500 | 10.9 | 6.6 | 19.8 | 21.8 | 5.0 | 1.0 | 2.9 | Yes |
|  | NIMR |  | Susceptible | 9.8 | 8.3 | 14.2 | 19.6 | 5.0 | 1.0 | 8.6 | Yes |
| Isocycloseram | IHI | Aedes<br>aegypti | Bagamoyo | 16.3 | 13.8 | 20.0 | 32.6 | 15.0 | 1.0 | 7.7 | Yes |
|  | IRD |  | Bora | 2.1 | 1.9 | 2.5 | 4.2 | >2.0 | 1.0 | 5.0 | Yes |
|  | LSTM |  | New Orleans | 2.7 | 2.3 | 3.4 | 5.5 | 2.0 | 1.0 | 6.6 | Yes |
|  | NIMR |  | Susceptible | 48.2 | 33.7 | 71.2 | 96.3 | 10.0 | 1.0 | 6.6 | Yes |
|  | CRID | Anopheles coluzzii | N'gousso | 1.2 | 1.0 | 1.9 | 2.3 | 1.0 | 1.0 | 2.0 | No† |
|  | CRID | Anopheles funestus | FANG | 28.2 | 13.5 | 71.8 | 56.5 | 10.0 | 0.9 | 11.1 | Yes |
|  | LSTM |  | FANG | 290.3 | 164.6 | 48940.2 | 580.6 | 100.0 | 0.6 | 17.7 | Yes |
|  | NICD |  | FANG | 26.1 | 22.2 | 33.1 | 52.2 | 25.0 | 1.0 | 4.9 | Yes |
|  | CRID | Anopheles gambiae | Kisumu | 1.2 | 1.0 | 2.1 | 2.4 | 1.0 | 1.0 | 3.8 | No† |
|  | IHI |  | Kisumu | 21.3 | 17.5 | 30.3 | 42.7 | 15.0 | 0.9 | 10.7 | Yes |
|  | IRD |  | Kisumu | 0.9 | 0.8 | 1.3 | 1.9 | 1.0 | 1.0 | 5.5 | Yes |
|  | LSTM |  | Kisumu | 7.1 | 6.2 | 8.5 | 14.2 | 6.0 | 1.0 | 5.5 | Yes |
|  | LSHTM | Anopheles stephensi | SDA500 | 0.9 | 0.6 | 2.1 | 1.9 | 2.5 | 1.0 | 0.1 | No† |
|  | LSTM |  | SDA500 | 16.9 | 12.8 | 24.8 | 33.8 | 25.0 | 1.0 | 5.5 | Yes |
|  | NIMR |  | Susceptible | 24.8 | 20.4 | 31.3 | 49.6 | 25.0 | 0.8 | 17.3 | Yes |

†Results marked as not converged should be interpreted with caution. For CRID Anopheles spp. (Broflanilide), data comprised only 1–2 replicates with mortality at or near 100% across all tested concentrations, meaning the dose–response curve could not be adequately resolved and LC99 estimates extrapolate below the lowest tested dose. For CRID Anopheles spp. (Isocycloseram) and LSHTM An. stephensi SDA500 (Isocycloseram), the tested concentration range was insufficient to capture the full dose–response relationship, precluding reliable convergence.

##### Step 3. Multi-centre validation of tentative discriminating concentrations against all mosquito species

Tentative discriminating concentrations (TDCs) were proposed for each testing centre for each compound against *An. gambiae, Ae. aegypti, An. funestus*, and *An. stephensi* based on data from that centre (Table 4). These values were determined using either a calculated 2×LC□□ or an “observed LC□□□.” The observed LC□□□ was used in cases where LC estimates differed substantially between centres and the highest calculated LC□□ was unrealistically high compared with the data generated elsewhere. In those situations, the observed LC□□□ was defined as the lowest tested concentration that consistently produced 100 percent mortality at the centre with the highest LC estimate, provided this value was not more than twofold higher than the highest confidence 2×LC□□ from the remaining sites. Each proposed TDC was tested by at least three independent laboratories to validate its suitability as a proposed DC. To address the large discrepancies observed in DC estimates from Step 2, IRD and IHI conducted additional Step 3 tests using additional doses. IRD tested *Ae. aegypti* with broflanilide at 3 µg and isocycloseram at 6 µg, and *An. gambiae* with broflanilide at 3 µg and isocycloseram at 2 µg. IHI also tested *Ae. aegypti* with Isocycloseram at 6 µg. These additional data were generated to support the selection of final DC values.

**Table 4.** Summary of mortality results from bottle bioassays conducted across multiple test centres using broflanilide- and isocycloseram-coated bottles. Each centre tested relevant mosquito species at control (0 µg/bottle) and selected validation concentrations, recording 24-hour post-exposure mortality. All data represent a single biological replicate except for NICD data where an additional two biological replicates were completed

| Centre | Species | Insecticide | Concentration (µg/bottle) | 24 Hour Mortality (%) |
| --- | --- | --- | --- | --- |
| CRID | *An. funestus* | Broflanilide | 0 | 8 |
|  |  |  | 14 | 100 |
|  |  | Isocycloseram | 0 | 8 |
|  |  |  | 57 | 100 |
|  | *An. gambiae* | Broflanilide | 0 | 18 |
|  |  |  | 11 | 100 |
|  |  | Isocycloseram | 0 | 18 |
|  |  |  | 30 | 100 |
| IHI | *Ae. aegypti* | Broflanilide | 0 | 0 |
|  |  |  | 10 | 100 |
|  |  | Isocycloseram | 0 | 0 |
|  |  |  | 6 | 100 |
|  |  |  | 0 | 0 |
|  |  |  | 15 | 100 |
|  | *An. gambiae* | Broflanilide | 0 | 0 |
|  |  |  | 11 | 100 |
|  |  | Isocycloseram | 0 | 0 |
|  |  |  | 30 | 100 |
| IRD | *Ae. aegypti* | Broflanilide | 0 | 0 |
|  |  |  | 3 | 100 |
|  |  |  | 10 | 100 |
|  |  | Isocycloseram | 0 | 0 |
|  |  |  | 6 | 100 |
|  |  |  | 15 | 100 |
|  | *An. gambiae* | Broflanilide | 0 | 1 |
|  |  |  | 3 | 100 |
|  |  |  | 11 | 100 |
|  |  | Isocycloseram | 0 | 1 |
|  |  |  | 2 | 100 |
|  |  |  | 30 | 100 |
| LSHTM | *An. stephensi* | Broflanilide | 0 | 2 |
|  |  |  | 22 | 100 |
|  |  | Isocycloseram | 0 | 0 |
|  |  |  | 50 | 100 |
| LSTM | *An. funestus* | Broflanilide | 0 | 2 |
|  |  |  | 14 | 100 |
|  |  | Isocycloseram | 0 | 0 |
|  |  |  | 57 | 100 |
|  | *An. gambiae* | Broflanilide | 0 | 0 |
|  |  |  | 11 | 100 |
|  |  | Isocycloseram | 0 | 2 |
|  |  |  | 30 | 100 |
|  | *An. stephensi* | Broflanilide | 0 | 2 |
|  |  |  | 22 | 100 |
|  |  | Isocycloseram | 0 | 0 |
|  |  |  | 50 | 100 |
| NIMR | *Ae. aegypti* | Broflanilide | 0 | 0 |
|  |  |  | 10 | 100 |
|  |  | Isocycloseram | 0 | 0 |
|  |  |  | 15 | 100 |
|  | *An. stephensi* | Broflanilide | 0 | 0 |
|  |  |  | 22 | 100 |
|  |  | Isocycloseram | 0 | 0 |
|  |  |  | 50 | 100 |
| NICD | *An. funestus* | Broflanilide | 0 | 0 |
|  |  |  | 14 | 99 |
|  |  |  | 15 | 100 |
|  |  | Isocycloseram | 0 | 0 |
|  |  |  | 57 | 93 |
|  |  |  | 60 | 100 |

Most Step 3 tests gave 100% mortality across all doses tested in all test centres, and so TDCs were confirmed. The only dataset where this was not the case was NICD data where 14µg broflanilide gave 98% mortality and 57µg isocycloseram gave 93% mortality for *An. funestus*. A second experiment was done at NICD with two technical replicates of 14µg broflanilide which gave 99% mortality. Finally, a set of 3 replicates was done using 15µg broflanilide and 60µg isocycloseram which gave 100% for both compounds across all three replicates. The DCs were therefore deemed to be appropriate, albeit with a caveat that variable results were observed at this one testing site.

The DCs based on the total data set from this study are shown in Table 5. Some values were rounded for simplicity to generate the final DCs recommended to WHO for adoption for each compound-species pair.

**Table 5.**
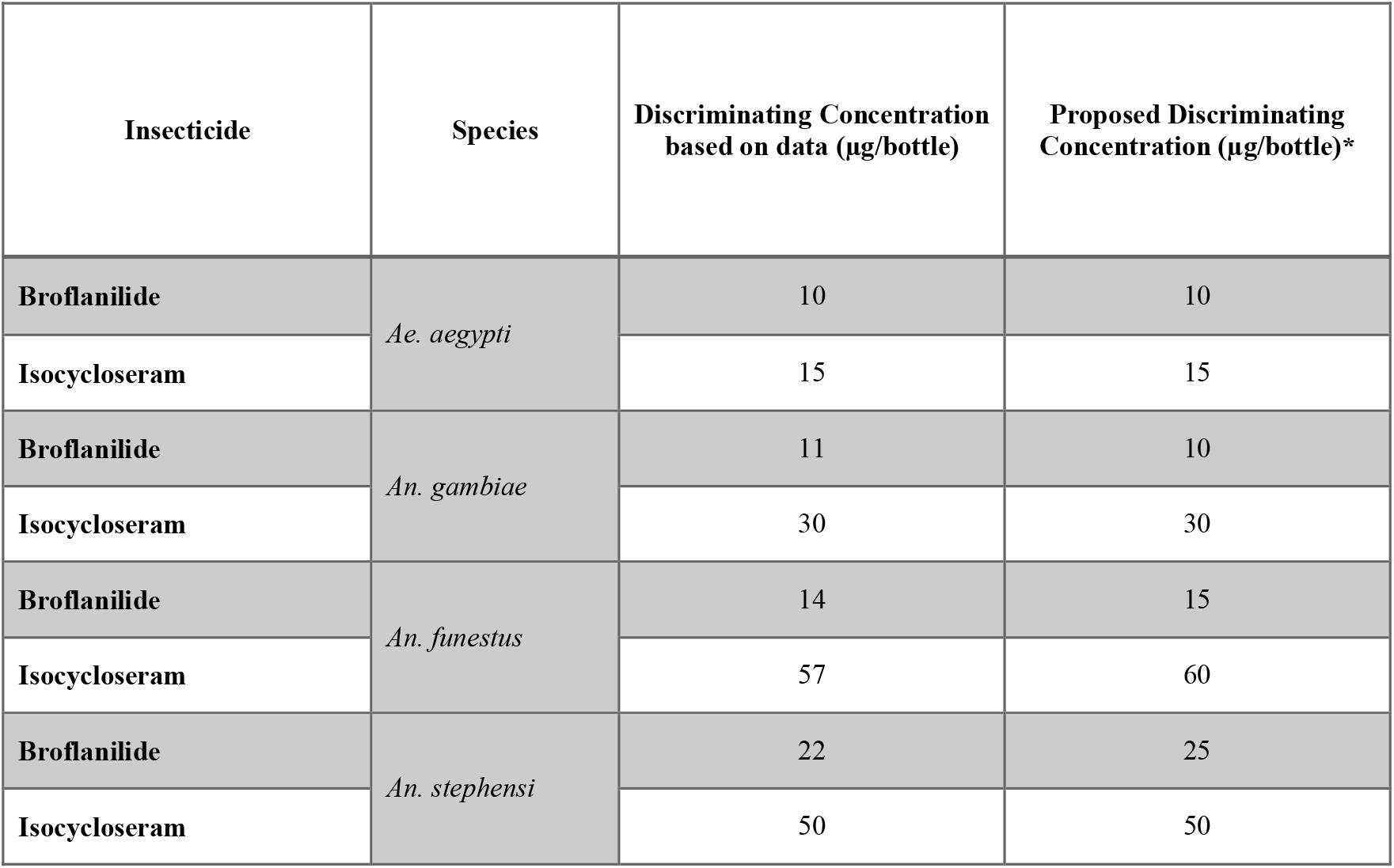
Proposed tentative discriminating concentrations (TDCs) for broflanilide and isocycloseram in WHO bottle bioassays against Aedes and Anopheles species based on the analysed Step 2 data.

| Insecticide | Species | Discriminating Concentration based on data (µg/bottle) | Proposed Discriminating Concentration (µg/bottle)* |
| --- | --- | --- | --- |
| Broflanilide | *Ae. aegypti* | 10 | 10 |
| Isocycloseram |  | 15 | 15 |
| Broflanilide | *An. gambiae* | 11 | 10 |
| Isocycloseram |  | 30 | 30 |
| Broflanilide | *An. funestus* | 14 | 15 |
| Isocycloseram |  | 57 | 60 |
| Broflanilide | *An. stephensi* | 22 | 25 |
| Isocycloseram |  | 50 | 50 |

## 5 Discussion

The WHO provides guidance on how to assess insecticide resistance in mosquito vectors in a standardised way using a discriminating concentration (DC) assay (1) designed to detect the early signs of emerging resistance to an insecticide. The data generated by monitoring mosquito populations for their susceptibility to a range of insecticides can inform decisions by those designing and implementing vector control strategies.

A WHO-sponsored multi-centre study was conducted in 2024–2025 to propose DCs for two new insecticides, broflanilide and isocycloseram, for resistance monitoring in *Ae. aegypti, An. Gambiae s*.*s*., *An. funestus*, and *An. stephensi*. Both insecticides belong to IRAC class 30 and are incorporated into vector control products which are prequalified by WHO for public health use. Seven laboratories from Africa, Asia, and Europe participated, coordinated by the Liverpool School of Tropical Medicine (LSTM) in collaboration with World Health Organization (WHO).

The study followed a standardised, three-step approach: preliminary testing to establish suitable concentration ranges (Step 1), multi-centre concentration–response testing to estimate LC_99_/LC_100_ and propose tentative discriminating concentrations (TDCs) (Step 2), and multi-centre validation of the TDCs (Step 3). Well-characterised, fully susceptible laboratory colonies were used for all assays. WHO bottle bioassays, with 1 h exposure, 24 h drying, and 24 h holding time, were confirmed as the most appropriate method for both compounds, with MERO surfactant at 1,500 ppm for *Aedes* spp. and 800 ppm for *Anopheles* spp.

In total, 62,673 mosquitoes were tested in WHO bottle assays across all centres. Validated TDCs were established for each species-compound combination, generally supported by three fully replicated datasets from partner sites, along with separate confirmation that the selected TDC killed 100% of all susceptible strains tested. Although generally successful in generating the desired dataset, the study was met with some logistical challenges which impacted the data that could be collected. Limited capacity for *An. stephensi* testing following the unforeseen withdrawal of one testing centre from the study left only LSTM able to test with this species. The London School of Hygiene and Tropical Medicine (LSHTM) held an independent colony of the same strain, SDA500, and although they were unable to join the study they provided mosquitoes to LSTM. Although this ensured *An. stephensi* testing could proceed in three sites, the lack of multi-centre data restricted inter-laboratory comparison and reduced the strength of evidence for validation. The delays caused by this change to the study plan meant that seven rather than the nine planned replicate tests were possible. The completion rate for all other insecticide-species combinations was 100%.

Some suggestions on the methods used to generate and analyse data for the purpose of establishing DCs are proposed based on the experience gained during this multi-centre study under four headings, below.

### 5.1.1 Selecting concentrations for Step 2

An LC_99_ is an inherently challenging target to estimate due to its dependence on observations very close to, but not reaching, 100% mortality. Mortality observations of 100% are far less useful for this purpose as they only identify the LC_99_ threshold has been exceeded, but not by how much. As a result, LC_99_ estimates carry substantial error unless the experimental design is explicitly targeted at observing the 95-99% range.

A common misconception is that a full dose response curve with a wide spread of mortality values is required for the purpose of obtaining the target LC value. For robust LC_99_ estimates, rich information near the target value matters far more than accurately defining the shape of the lower parts of the curve. Thus, to estimate a target LC value it is ideal to conduct a preliminary range finding step that approximates the concentration needed, followed by a targeted step that aims above and below that dose range. As highlighted previously (9,10), more robust bioassay data does not necessarily require larger sample sizes as instead mosquitoes can be better targeted towards relevant concentrations. In the final analysis, data generated during the range finding step can be included in the final modelling analysis to strengthen the estimates.

### 5.1.2 Ensuring power for robust LC calculations

Sample size calculations should focus on the precision required for the LC estimate. In principle the first decision is the level of error that can be tolerated, for example ±10 percent. In practice the process becomes a comparison of alternative sample size scenarios and the resource implications of each. Precision depends on the number of concentrations tested and the number of replicates at each concentration. For any fixed total number of mosquitoes there will be an optimal balance between these two elements. Identifying the minimum total number of mosquitoes for a given level of precision is an optimisation problem that usually requires simulation. Such approaches are well established but rarely straightforward for end users because they rely on realistic assumptions about variance that must be informed by existing data or set conservatively (8).

Although the LC_99_ is conceptually the most relevant value for setting a discriminating concentration, its practical limitations outlined above and in (7), and resource limitations mean that a trade-off exists between the usefulness of the LC chosen and the robustness of its estimate. An LC_50_ is typically estimated with the greatest precision because data are generally more balanced above and below that point (unless explicitly designed otherwise). Intermediate values such as LC_80_ or LC_90_ may offer a more realistic compromise between biological relevance and statistical confidence.

### 5.1.3 Defining a DC

While the statistical principles underlying estimation of LC values are well defined, the step of converting an LC to a discriminating concentration is open to improvement. Notably, the confidence interval around the LC estimate is not currently considered in the calculation of the final DC (3). Even when using a Bayesian modelling approach that better captures heterogeneity, the uncertainty interval is not taken into account in this final step (14). If the true LC_99_ lies at the lower bound of the confidence interval, a discriminating concentration based on the mean could be set too high, reducing the sensitivity of the assay, and early signs of resistance may be missed. Thus, a conservative approach would be to use the lower bound of the 95 percent confidence interval when defining the discriminating concentration.

The current practice of doubling the LC_99_ assumes that unobserved variation in susceptible populations needs to be accommodated, i.e. the range of susceptible strains assessed does not capture the range of all possible susceptible populations. However, the choice of a multiplier of two appears arbitrary. If the multiplier is set too high there is a risk that the early sign of resistance will be missed.

Increasing the robustness of LC estimates could reduce or remove the need for such a multiplier if the susceptible populations used were reasonably representative. In recognition of this, the study design applied here and proposed by WHO (3) tests three susceptible strains of each species and adopts the highest LC_99_ or LC_100_ as the basis for the DC. The more strains which are included, the greater the confidence that the full range of susceptible phenotypes is represented, but in practice a multiplier may still be required, depending on the selected LC value.

#### 4. Setting DC values

Deciding on a DC is a challenge in situations in which LC estimates vary substantially between testing centres or susceptible strains. Selecting the highest value is sensible provided there is no evidence of quality control issues affecting one or more strains e.g. genetic contamination causing resistance, or poor rearing leading to smaller adults. All strains should be profiled (11) for phenotypic susceptibility before testing, but additional checks may be needed when results appear anomalous. These could include genotyping for known resistance markers or morphometric comparisons such as wing measurements. If no explanation for an outlier can be found, all data should remain in scope, and the highest LC should be used nonetheless to avoid setting a DC too low and thus causing false positive reports of resistance. If resources allow, further strains or repeated testing could help determine whether the outlier represents true biological variation or an anomaly. Even so, the decision needs to be made whether to retain the highest LC or exclude problematic data and this decision should be guided by clear criteria.

## Conclusion

This multi-centre study is a validation of the generic protocol for establishing DCs published by WHO (3), having successfully generated and validated discriminating concentrations for broflanilide and isocycloseram against An. gambiae, An. funestus, An. stephensi, and Ae. aegypti. This is the first time a DC has been established for isocycloseram. For broflanilide the DC’s generated reflect the DCs previously proposed, including the large variation between different sites with susceptible mosquitoes of the same species (12,13). These values will enable national programmes and research institutions to integrate these insecticides into their resistance monitoring activities.

## Declarations

### Author Contributions

Giorgio Praulins – Conceptualization, Investigation, Methodology, Data Curation, Formal Analysis, Visualization, Writing – Original Draft Preparation, Writing – Review & Editing, Project Administration.

Frank Mechan – Formal Analysis, Methodology, Visualisation, Writing – Review & Editing

Gemma Harvey - Formal Analysis, Methodology, Visualisation, Writing – Review & Editing

Basil Brooke - Investigation, Writing – Review & Editing, Supervision

Vincent Corbel - Investigation, Writing – Review & Editing, Supervision

Stéphane Duchon - Investigation, Writing – Review & Editing

Maria Kaiser - Investigation, Writing – Review & Editing

Sarah Moore - Investigation, Writing – Review & Editing, Supervision

Ahmadi Bakari Mpelepele - Investigation, Writing – Review & Editing

Shüné Oliver - Investigation, Writing – Review & Editing

Himmat Singh - Investigation, Writing – Review & Editing, Supervision

Jennifer Stevenson - Investigation, Writing – Review & Editing, Supervision

Yvan G. Fotso Toguem - Investigation, Writing – Review & Editing

Vaishali Verma - Investigation, Writing – Review & Editing

Charles Wondji - Investigation, Writing – Review & Editing, Supervision

Rosemary Susan Lees – Conceptualization, Funding Acquisition, Supervision, Writing – Original

Draft Preparation, Writing – Review & Editing, Project Administration

### Ethical approval and consent

Ethical approval and consent were not required.

## Acknowledgments

We thank Mitsui Chemicals Crop & Life Solutions, Inc. for provision of broflanilide for the study, and Syngenta Crop Protection AG for provision of isocycloseram. We are extremely grateful to the project sponsors, World Health Organization (WHO), and particularly Seth Irish and Emmanuel Chanda for technical guidance and consultation throughout the study. We offer great thanks to Dr Mojca Kristan at the London School of Hygiene & Tropical Medicine (LSHTM) for provision of mosquitoes for this study in replacement of a testing centre that had to drop out of the study at the last minute.

## Data Availability Statement

The datasets generated for this study can be found in the Zenodo repositors at https://zenodo.org/records/17804435.

