## Supplementary Material for "Multi-centre laboratory study to determine discriminating concentrations for broflanilide and isocycloseram resistance monitoring in mosquitoes"

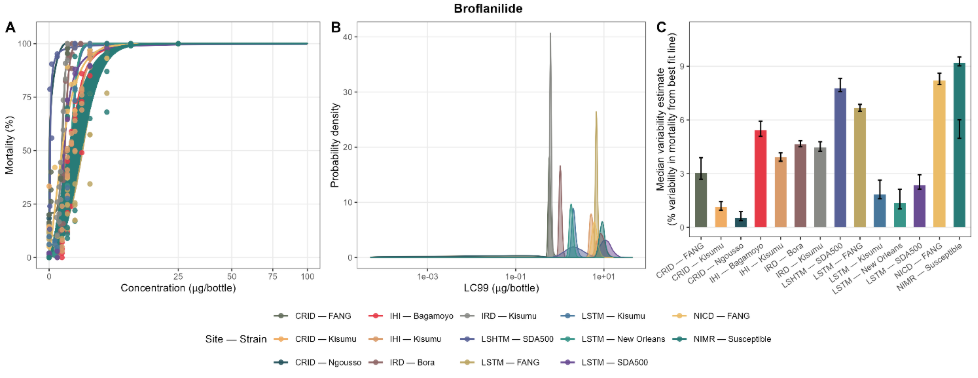

Figure S1. Concentration–response relationships, LC_99_ posterior distributions, and within-bioassay variability for (A–C) broflanilide across all participating sites and mosquito strains in WHO bottle bioassays. Panel A shows fitted concentration–mortality curves with 95% confidence intervals (shaded), with observed data points overlaid; x-axes are on a square-root scale. Panel B shows the posterior probability density of LC_99_ estimates for each site–strain combination within the plausible concentration range; x-axes are on a log scale. Panel C shows the median within-bioassay variability estimate (mean absolute percentage deviation from the best-fit line across all MCMC iterations) with 95% confidence intervals. In all panels, lines, shading, and points are coloured by site–strain combination as shown in the legend. Site–strain combinations where LC_99_ could not be reliably estimated within the tested concentration range are excluded from panel B.

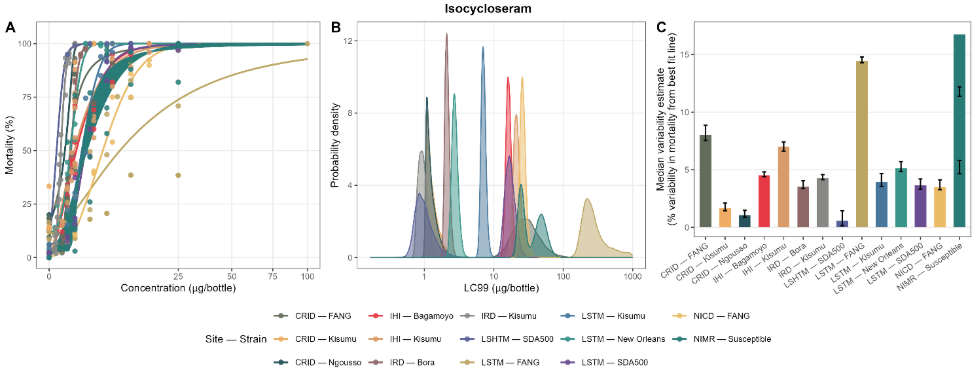

Figure 11. Concentration–response relationships, LC_99_ posterior distributions, and within-bioassay variability for (A–C) isocycloseram across all participating sites and mosquito strains in WHO bottle bioassays. Panel A shows fitted concentration–mortality curves with 95% confidence intervals (shaded), with observed data points overlaid; x-axes are on a square-root scale. Panel B shows the posterior probability density of LC_99_ estimates for each site–strain combination within the plausible concentration range; x-axes are on a log scale. Panel C shows the median within-bioassay variability estimate (mean absolute percentage deviation from the best-fit line across all MCMC iterations) with 95% confidence intervals. In all panels, lines, shading, and points are coloured by site–strain combination as shown in the legend. Site–strain combinations where LC_99_ could not be reliably estimated within the tested concentration range are excluded from panel B.

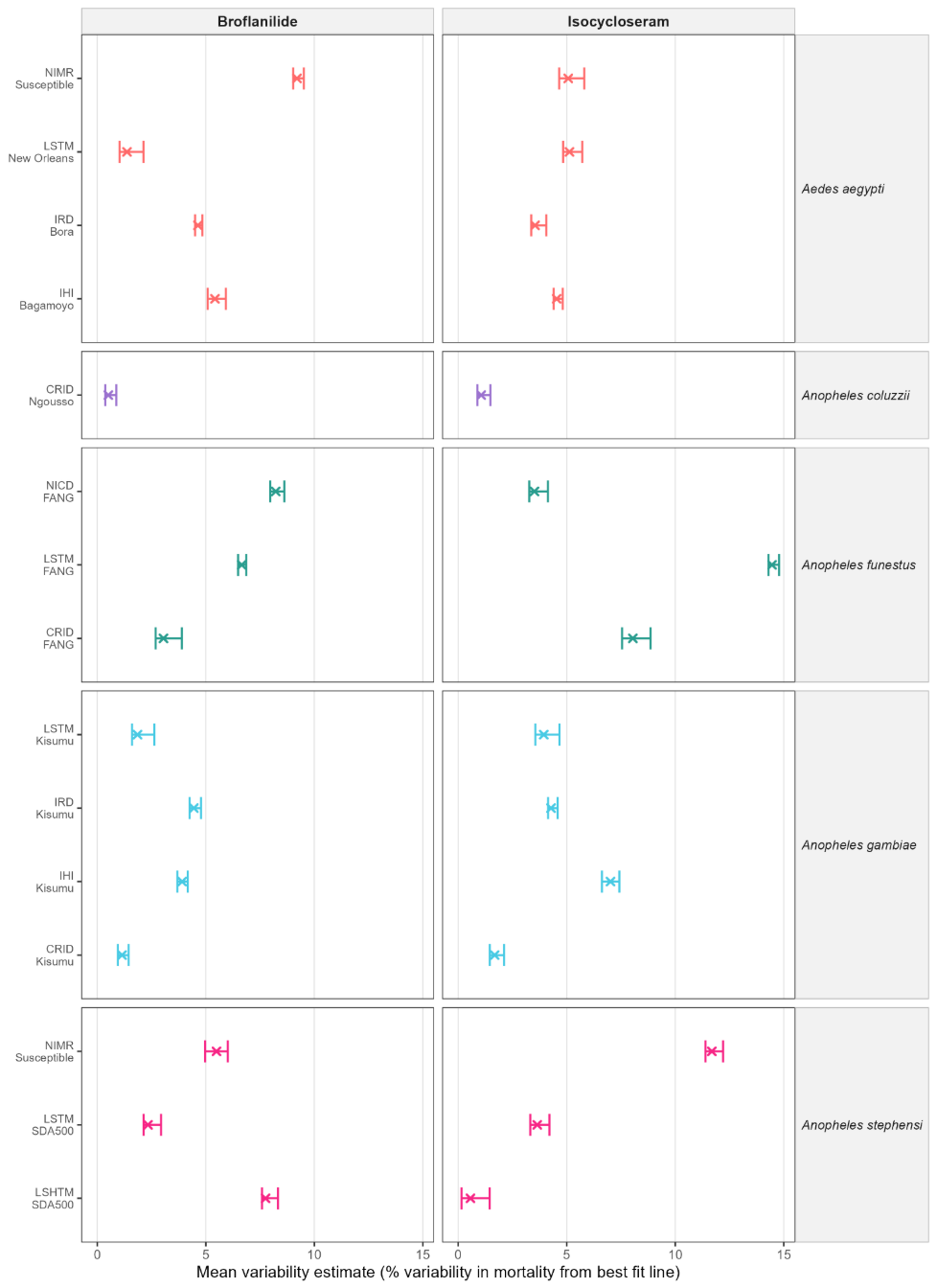

Figure 12. Within-bioassay variability in mortality estimates for broflanilide and isocycloseram across all participating sites and mosquito strains. Points (×) represent the median variability estimate, expressed as the mean absolute percentage deviation of individual data points from the best-fit concentration–response line, computed across all Markov Chain Monte Carlo iterations. Horizontal error bars show the 95% confidence intervals. Panels are faceted by mosquito species (rows) and insecticide (columns). Points are coloured by species. Results are shown for all site–strain combinations where model convergence was achieved.

Table **S1**. Summary of additional statistics for broflanilide and isocycloseram across all participating centres, mosquito species, and strains in WHO bottle bioassays.

| **Insecticide** | **Site** | **Species** | **Strain** | **Intercept** | **Intercept SE** | **Intercept p** | **Slope** | **Slope SE** |
| --- | --- | --- | --- | --- | --- | --- | --- | --- |
| **Broflanilide** | IHI | *Aedes aegypti* | Bagamoyo | 0.3542 | 2.5789 | 0.8914 | 0.9924 | 0.0382 |
| **Broflanilide** | IRD | *Aedes aegypti* | Bora | -0.1349 | 2.4883 | 0.9572 | 1.0022 | 0.0367 |
| **Broflanilide** | LSTM | *Aedes aegypti* | New Orleans | -0.3089 | 0.6356 | 0.6357 | 1.0034 | 0.0102 |
| **Broflanilide** | NIMR | *Aedes aegypti* | Susceptible | 0.1108 | 8.1211 | 0.9893 | 0.9971 | 0.1033 |
| **Broflanilide** | CRID | *Anopheles coluzzii* | N’gousso | 1.3219 | 0.911 | 0.1631 | 0.9868 | 0.0098 |
| **Broflanilide** | CRID | *Anopheles funestus* | FANG | 1.3287 | 2.254 | 0.561 | 0.9865 | 0.0292 |
| **Broflanilide** | LSTM | *Anopheles funestus* | FANG | 0.0095 | 2.8828 | 0.9974 | 0.9981 | 0.043 |
| **Broflanilide** | NICD | *Anopheles funestus* | FANG | -0.1299 | 6.1006 | 0.9832 | 0.9999 | 0.0806 |
| **Broflanilide** | CRID | *Anopheles gambiae* | Kisumu | 2.5331 | 2.5845 | 0.3394 | 0.9747 | 0.0278 |
| **Broflanilide** | IHI | *Anopheles gambiae* | Kisumu | 0.0431 | 1.5619 | 0.9781 | 0.9996 | 0.0234 |
| **Broflanilide** | IRD | *Anopheles gambiae* | Kisumu | -0.052 | 2.0311 | 0.9798 | 0.9972 | 0.0331 |
| **Broflanilide** | LSTM | *Anopheles gambiae* | Kisumu | 0.1001 | 0.936 | 0.9166 | 1.0002 | 0.0152 |
| **Broflanilide** | LSHTM | *Anopheles stephensi* | SDA500 | 1.469 | 7.3942 | 0.8458 | 0.9775 | 0.1018 |
| **Broflanilide** | LSTM | *Anopheles stephensi* | SDA500 | 0.0008 | 1.5659 | 0.9996 | 0.9988 | 0.0206 |
| **Broflanilide** | NIMR | *Anopheles stephensi* | Susceptible | -0.163 | 3.588 | 0.9642 | 1.0034 | 0.0512 |
| **Isocycloseram** | IHI | *Aedes aegypti* | Bagamoyo | 0.2889 | 2.392 | 0.9046 | 0.9952 | 0.0366 |
| **Isocycloseram** | IRD | *Aedes aegypti* | Bora | -0.4157 | 1.9605 | 0.8343 | 1.0072 | 0.0308 |
| **Isocycloseram** | LSTM | *Aedes aegypti* | New Orleans | -2.0496 | 3.6946 | 0.5893 | 1.019 | 0.0518 |
| **Isocycloseram** | NIMR | *Aedes aegypti* | Susceptible | -0.9855 | 2.5179 | 0.6998 | 1.0216 | 0.0379 |
| **Isocycloseram** | CRID | *Anopheles coluzzii* | N’gousso | 1.4256 | 1.4247 | 0.3296 | 0.9853 | 0.0154 |
| **Isocycloseram** | CRID | *Anopheles funestus* | FANG | 0.0466 | 4.6695 | 0.9922 | 0.9968 | 0.0774 |
| **Isocycloseram** | LSTM | *Anopheles funestus* | FANG | 0.7286 | 6.1513 | 0.9065 | 0.9804 | 0.1402 |
| **Isocycloseram** | NICD | *Anopheles funestus* | FANG | 0.1338 | 2.5428 | 0.9589 | 0.9965 | 0.0384 |
| **Isocycloseram** | CRID | *Anopheles gambiae* | Kisumu | 2.2485 | 2.8165 | 0.4346 | 0.9771 | 0.0305 |
| **Isocycloseram** | IHI | *Anopheles gambiae* | Kisumu | 0.0831 | 3.449 | 0.9809 | 0.9991 | 0.0529 |
| **Isocycloseram** | IRD | *Anopheles gambiae* | Kisumu | 0.025 | 1.6798 | 0.9882 | 1.0025 | 0.0292 |
| **Isocycloseram** | LSTM | *Anopheles gambiae* | Kisumu | -0.3805 | 2.9569 | 0.8997 | 1.0047 | 0.0433 |
| **Isocycloseram** | LSHTM | *Anopheles stephensi* | SDA500 | -1.3998 | 0.1136 | 0.0001 | 1.0135 | 0.0012 |
| **Isocycloseram** | LSTM | *Anopheles stephensi* | SDA500 | -0.3175 | 2.1312 | 0.8831 | 1.0027 | 0.0319 |
| **Isocycloseram** | NIMR | *Anopheles stephensi* | Susceptible | 0.4375 | 7.2829 | 0.9527 | 0.9879 | 0.1153 |

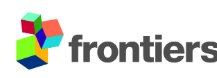
